# Ingestion of poisonous eggs from the invasive apple snail *Pomacea canaliculata* adversely affects bullfrog *Lithobathes catesbeianus* intestine morphophysiology

**DOI:** 10.1101/2020.07.07.191684

**Authors:** Tabata R. Brola, Marcos S. Dreon, Patricia E. Fernández, Enrique L. Portiansky, Horacio Heras

## Abstract

The poisonous eggs of *Pomacea canaliculata* (Caenogastropoda, Ampullariidae) hardly have any predator. This invasive snail, listed among the 100 worst invasive species, is a serious crop pest and a vector of a human parasitosis. Females lay eggs in pink-reddish masses, presumably as a warning coloration of their chemical defenses. Egg ingestion alters rodents’ gastrointestinal tract and is lethal if injected, but its effect on other taxa is unknown. Here we explored the toxic effects of *Pomacea canaliculata* eggs on bullfrog *Lithobathes catesbeianus* (Anura, Ranidae).

Juvenile bullfrogs were gavaged with egg extracts and their digestive tract analyzed after 24 h and 48 h using histological, immunohistochemical and lectin-histochemical techniques. Toxicity was also evaluated by intraperitoneal injection of eggs extracts.

Egg extract ingestion adversely affected small intestine of bullfrogs. Short term (24 h) effects included large, reversible changes of the intestinal wall, villi morphology, and changes in the glycosylation patterns of enterocytes. The mucosal area increased and infiltration of inflammatory cells, mainly eosinophils and macrophages, were observed together with a weak hemorrhage. Most of these changes reversed after 48 h. Besides, intraperitoneal injection of egg extract was nontoxic to bullfrog juveniles and no lethality or behavioral changes were observed, a remarkable difference with egg’ effect on mammals.

As a whole, these results indicate that toxins of apple snail eggs reversibly modify gut morphology, which may alter bullfrog physiology limiting their ability to absorb egg nutrients. This study extends the known targets of the apple snail egg defenses against predation to amphibians.

## INTRODUCTION

Eggs of most animals are subject to strong predation (Ricklefs 1969, Kamler 2005). One of the exemptions are the eggs of the freshwater apple snail *Pomacea canaliculata* (Lamarck, 1822) (Caenograstropoda, Ampullariidae). Native from South America, this snail has been introduced worldwide becoming an invasive species and a vector of a human parasitosis (Hayes et al. 2008; Lv et al. 2011). Their invasiveness is partly related to their reproductive strategy (Lowe et al. 2000; Martin and Estebenet 2002, Hayes et al. 2012), especially to their egg defenses (Heras et al. 2008, Dreon et al. 2010, 2013, 2014). Females deposit pink-reddish egg masses outside the water on a weekly basis, each containing 30–300 eggs (Albrecht et al. 1996; Cadierno et al. 2018) filled with a perivitelline fluid (PVF) that surrounds the developing embryo. PVF is known to serve the critical functions of embryonic nutrition, protection and defense in *P. canaliculata* (Heras et al. 1998; Heras et al. 2007). Thus, the PVF enable eggs to develop under harsh aerial conditions including desiccation high temperatures, and help coping with terrestrial predators (Dreon et al. 2004; Heras et al. 2007; Dreon et al. 2007). Even more, predators that feed on adults, like the snail kite *Rostrhamus sociabilis* (Vieillot, 1817) or the Norway rat *Rattus norvegicus* (Berkenhout, 1769), systematically discard the albumen gland where most egg PVF is synthetized (Przeslawski 2004, Cadierno et al. 2017b,a, Hayes et al. 2015).

The PVF of *P. canaliculata* contains more than 30 proteins (perivitellins) (Sun et al. 2012; Sun et al. 2019) including a hyperstable non-digestible carotenoprotein (PcOvo), massively accumulated in the egg. PcOvo has a dual function, on the one hand it can resist proteolysis and withstands passage through the digestive tract of rodents almost unaffected, lowering the nutritional value of the eggs (Pasquevich et al. 2017; Dreon et al. 2010; Dreon et al. 2014). On the other hand, it provides clutches with a vivid coloration (aposematic), probably a warning to visual hunting predators of their toxic defenses. The PVF defensive cocktail also includes a neuro- and enterotoxic lectin (PcPV2), lethal to mice when intraperitoneally injected, that adversely affects their gastrointestinal tract if ingested (Heras et al. 2008, Giglio et al. 2020). Furthermore, oral administration of *P. canaliculata* PVF to rodents alters their gut morphophysiology and decreases rat growth rate (Dreon et al. 2014; Giglio et al. 2018). These unusually efficient biochemical defenses may explain the lack of reported predators for the eggs (Cadierno et al. 2017a). However, most of the studies on the defensive system of *P. canaliculata* eggs have been conducted using only mammals as models, and whether the ingestion of these perivitellins affect other taxa is unknown. Amphibians are a likely target of these defenses and their digestive physiology may differ considerably from that of mammals (Sabat and Bozinovic 2000). Besides, many anurans and urodeles overlap in their geographical distribution with those of the apple snail. In particular, the invasive bullfrog *Lithobathes catesbeianus* (Shaw, 1802) (Anura,Ranidae) partially overlaps with *P. canaliculata* geographical distribution in both their native and introduced regions (Akmentins and Cardozo 2010).

To explore possible targets of *P. canaliculata* egg defense system beyond mammals, and to further understand the biological-ecological meaning of the digestive plasticity in amphibians, we selected *L. catesbeianus* as a model of potential predator of the eggs. This North American frog, introduced to several parts of the world including South America, has voracious habits and a varied diet including eggs of both vertebrates and invertebrates species (Hirai 2004; Snow and Witmer 2010).

We report here that oral administration of *P. canaliculata* PVF to juvenile bullfrogs has a strong adverse effect on gut at tissue and organ levels, temporarily affecting their small intestine morphology and inducing an extended cellular reactive reaction. Most of these changes were reversed after 48 h. However, unlike mammals, no lethal or neurotoxic effects were observed when injected with snail egg extracts. As far as we know, the occurrence of toxins that affect intestinal morphology probably affecting absorption, has not been recognized among amphibians before. This is the first report of a poisonous effect by the ingestion of *P. canaliculata* egg extracts on a vertebrate other than mammals.

## METHODS

### Egg masses and PVF preparation

*P. canaliculata* egg clutches were collected from the field in La Plata city, Buenos Aires, Argentina, between November and March during the reproductive season. Only eggs not beyond morula stage were used. PVF was prepared following Dreon et al. (2014). Briefly, eggs were homogenized in ice cold 20 mM Tris-HCl, pH 7.5 in a buffer: sample ratio of 3:1 v/w. The homogenate was sequentially centrifuged at 10,000 g for 30 min and at 100,000 g for 60 min, and the pellet discarded. The supernatant (egg PVF) was equilibrated in 50 mM phosphate buffer saline (PBS), pH 7.4 using a centrifugal filter device of 50 kDa molecular weight cut off (Millipore Corporation, MA). Total protein concentration was measured by the method of Lowry et al. 1951.

### Experimental animals

Sexually immature juvenile bullfrogs *Lithobathes catesbeianus* were purchased from an aquarium in Buenos Aires, Argentina, and bred in a specially conditioned room at the facilities of the School of Medicine, National University of La Plata. They were housed in plastic 60×40×50 cm containers with natural vegetation and soil. Water provided in plastic containers was changed daily. Frogs were maintained with a 12-h dark/light cycleat 22 ± 3 °C and fed once a day either with *Tenebrio* larvae or adult *Blaptica dubia*.

### Lethality test

To assess the PVF lethality on frogs, a preliminary screening test was performed. Groups of five bullfrog (4.0 ± 1.1 g) were intraperitoneally (i.p.) injected with a single dose of 200 μl of either PBS (control) or the same volume of a serial dilution of five concentrations of PVF in the range 1.5 – 12.0 mg/Kg using a 30G x 13 mm needle. As lethality was not observed, a second test was performed injecting three bullfrogs with 200 μl of 170 mg/Kg PVF, the highest possible concentration, which was 50 times higher than the LD_50_ established for mice (Heras et al. 2008). Animals were observed during 96 h after administration for lethality and behavioral changes.

### Histology analysis

Bullfrogs (n=6) were gavaged with a single dose of 200 μl of PVF (9 mg/ml) using a flexible plastic cannula. Control animals (n=3) were administrated with the same volume of PBS. Groups of three animals were euthanized either after 24 h or 48 h post-administration by immersion in a water bath containing Tricaine methane-sulfonate (MS-222) 0.2 mg/ml, following the procedure described by Tyler (2009). Intestines were removed and samples of the first part of the small intestine were cut, rinsed several times with PBS and then fixed in 10% neutral buffered formalin for 48 h for histological examination. After 48 h fixation, samples were transferred into 70° ethanol and dehydrated before being embedded in paraffin wax at 60°C for 1 h. Five to seven μm thick sections were obtained from the paraffin blocks using a manual microtome and routinely stained with haematoxylin and eosin (HE) according to Bancroft and Gamble (2008). Analysis was performed using an Olympus BX51 microscope. Digital images were obtained using an Olympus DP-70 video camera and then captured with an image analyzer (ImagePro Plus v.6.3, Media Cybernetics, USA).

### Morphometric analysis

Changes in small intestine morphometry caused by PVF administration were measured from images (4x magnification) obtained using the image analyzer (ImagePro Plus) from 9 properly oriented HE stained sections per animal, a total of 27 images for each experimental group. “Section area” (SA) was measured considering both mucosa (epithelium, and lamina propria) and the lumen area, but excluding muscular layer. “Tissue area” (TA) was measured considering the relative area of the mucosa to SA and excluding lumen area. The relation between lumen area and SA was also measured. To estimate the absorptive surface, perimeter of the epithelium facing lumen was measured in relation to SA.

### Presence of eosinophils and macrophages cells

A total of 27 small intestine sections for each group were analyzed to determine the presence of eosinophils and macrophages, which were counted in 5 randomly chosen fields of 0.05 mm^2^ from each section (60x magnification, 135 fields per experimental group). The presence of eosinophils was determined in HE stained sections. Macrophages were determined by immunohistochemistry (IHC). Briefly, sections were deparaffinized, hydrated through graded ethanol and incubated overnight at 4°C with a mouse anti-human CD68 clone KP1 monoclonal antibody (1:100, Dako, Denmark). Then, sections were revealed with a commercial kit (LSAB System-HRP, Dako, Denmark) that includes a streptavidin-biotin labeled secondary antibody. The colored reaction was developed using 3,3’-diaminobenzidine tetrahydrochloride (DAB, Dako, Denmark) (chromogen) and H_2_O_2_ as reaction substrates. All sections were then counterstained with Mayer’s hematoxylin before analysis. Dark brown stained cells were considered positive.

### Binding assay of PcPV2 toxic lectin to gut

The binding and location of PcPV2 to epithelial cells of the small intestine was performed on control, 24 h and 48 h post-gavage with PVF as described in Histology analysis section. Nine sections for each treatment were mounted on positively charged slides, deparaffinized, dehydrated, and incubated in 3% H_2_O_2_ in methanol for 30 min at room temperature. Slides were rinsed with PBS (pH = 7.4) and then subjected to antigen retrieval procedure using proteinase K 1:30 (Dako, Denmark). Nonspecific binding sites were blocked with 2% BSA for 30 min in a humid chamber at 4°C, followed by overnight primary anti-PcPV2 polyclonal antibody (1:200) incubation. Slides were then revealed using the Envision plus kit (Dako, Carpinteria, CA, USA). Polyclonal antibodies against PcPV2 were prepared in rabbits as previously described (Dreon et al. 2002). Positively immunostained regions showed a golden dark brown color using 3,3’-diaminobenzidine tetrahydrochloride (DAB) as chromogen and H_2_O_2_ reaction substrates (DakoCytomation, Glostrup, Denmark). All sections were counterstained with Maeyer’s hematoxylin.

### Collagen determination

Five 5 μm thick sections of the intestine of bullfrogs were stained using the Picrosirius red technique (Direct Red 80, Aldrich, Milwaukee, WI 53233, USA) for collagen evaluation, as described elsewhere (Portiansky et al. 2002). Briefly, sections were deparaffinized, hydrated through graded ethanol and stained for 1 h in a 0.1% solution of Sirius Red dissolved in aqueous saturated picric acid. Sections were then rapidly washed in running tap water and counterstained with Harris hematoxylin.

Samples observed under polarized light, using an analyzer (U-ANT, Olympus) and a polarizer (UPOT, Olympus) were used to study the birefringence of the stained collagen. Type I collagen reflected red to yellow light, while type III collagen reflected green light. Histological (20× magnification) images were digitized using a digital video camera (Olympus DP73, Japan) mounted on a widefield microscope (Olympus BX53, Japan). Captured images were saved in TIF format using an image analysis software (Olympus cellSens Dimension V1.7, Japan) automatically coordinated with the camera for later analysis. Total collagen was calculated as the sum of all connective tissue areas of the coronal sections (type I and type III collagen), divided by the total surface of the section.

### Lectinhistochemistry

To analyze possible changes in the glycosylation pattern of the enterocytes, tissues were evaluated with seven lectins (Lectin Biotinylated BK 1000 Kit, Vector Laboratories Inc., Carpinteria, CA, USA) namely: Con A (*Concanavalia ensiformis*), DBA (*Dolichos biflorus*), SBA (*Glycine max*), PNA (*Arachis hypogaea*), RCA-I (*Ricinus communis-I*), UEA-I (*Ulex europaeus-I*), and WGA (*Triticum vulgaris*). Briefly, 9 small intestine sections were deparaffinized with xylene dehydrated with 100% alcohol twice, 10 min each, and then endogenous peroxidase activity was quenched by incubating 5 min with 0.3-3.0% hydrogen peroxide in methanol. They were then hydrated, washed with PBS, and incubated with biotinylated lectins overnight. Sections were then washed with PBS, followed by 10-min incubation with streptavidin-HRP (streptavidin conjugated to horseradish peroxidase in PBS containing stabilizing protein and anti-microbial agents, Vector Laboratories Inc., USA). Finally, the bounded lectins were visualized incubating for 4-10 min with a buffered Tris-HCl solution (0.05 M, pH =6.0) containing 0.02% 3,3’-diaminobenzidine tetrahydrochloride (DAB) and 0.05% H_2_O_2_ (DAB; Dako, Carpinteria, USA). Sections were counterstain with Maeyer’s haematoxilyn. Enterocytes were divided into three zones for analysis: apical zone, stroma and supranuclear zone. Goblet cells were also analyzed. Tissues showing golden brown coloration were considered positive and given a qualitative value according to the intensity of color.

### Statistical analysis

A Kruskall-Wallis test was used to test morphometric differences. Differences between control, 24 h and 48 h post-gavage groups were determined by Dunn’s multiple comparison test. For absorptive surface analysis and collagen, one-way ANOVA was performed among control, 24 h and 48 h post-gavage groups. For post hoc comparison, the Tukey’s multiple comparison test was used. Statistical analysis was performed using Prism 5.03 (Graphpad Software Inc.). Results were considered significant at the 5% level.

## RESULTS

### Ingestion of egg PVF affects small intestine morphology of bullfrog

To evaluate the effect of gavage administration of PVF, the morphometry of bullfrog small intestine was analysed (Fig.1A). We found that after 24 h the small intestine mucosa area increased its area fraction with a mean of 0.80 ±0.05 as compared with the control group (0.62 ±0.08), while at 48h post-gavage the area fraction was reduced to 0.69±0.12 (Fig.1B). Simultaneously, lumen area fraction was significantly reduced after 24 h (0.20 ±0.08) as compared to controls (0.36±0.08), while at 48 h after gavage values were similar to those of controls (0.31±0.09) (Fig.1C). No changes in the absorptive surface were observed (Fig.1D). The villi of control animals were long with a continuous epithelium containing some goblet cells and well-defined central lymphatic capillaries (lacteal vessel) (Fig. 1E). After 24 h of PVF gavage, villi became wider almost obstructing lumen space, with the tip of the villi showing epithelial disorganization. Besides, the lamina propria was remarkably thickened (Fig.1F). The whole epithelium became discontinuous, with fewer goblet cells and a weak haemorrhage was observed (Fig.2). Forty-eight hours after gavage, the mucosa began to return to its normal morphology, showing narrower villi and wider lumen space than 24 h before. Nevertheless, the lamina propria was thicker than in controls (Fig. 1G).

**Figure 1.**
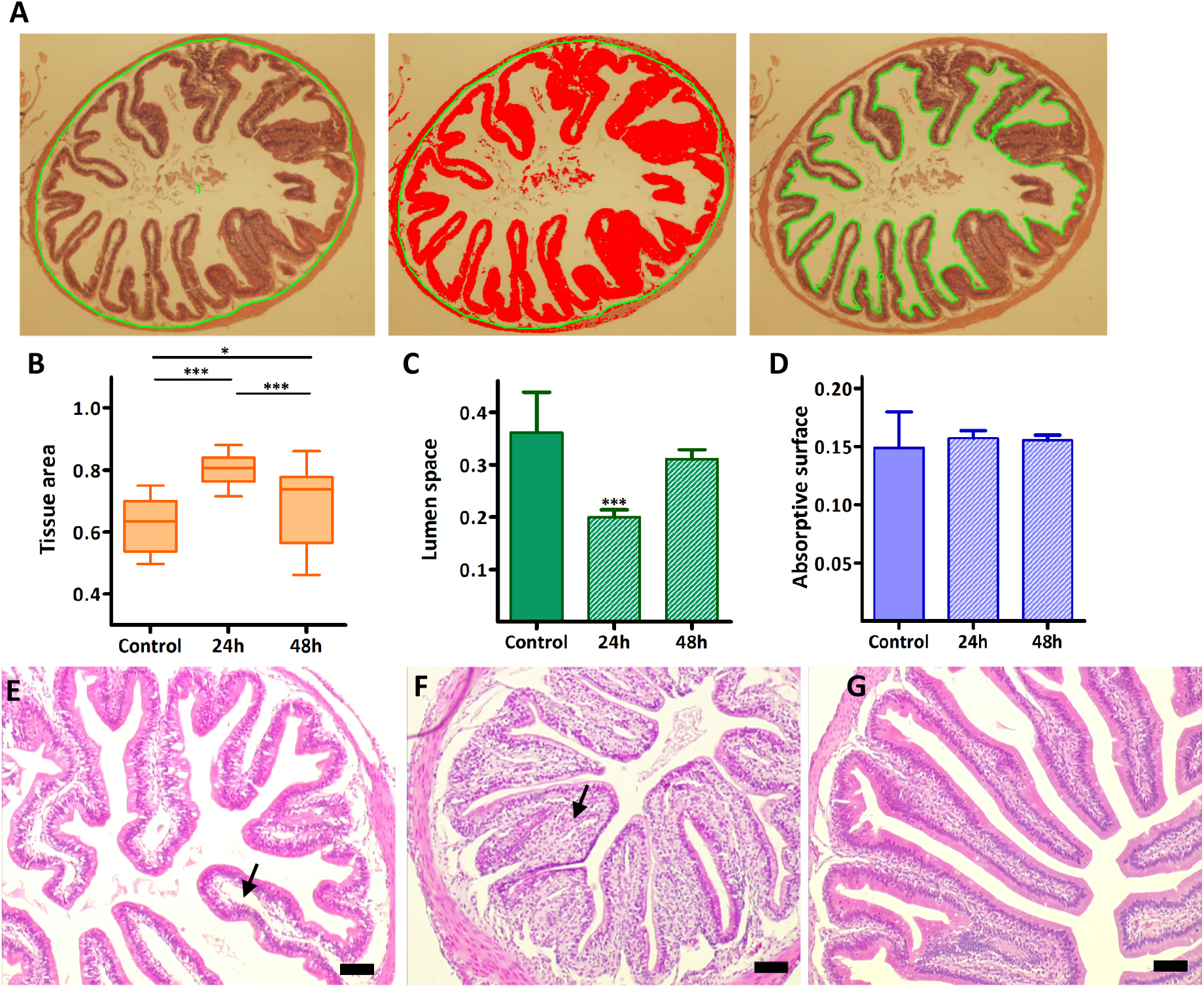
Changes in bullfrog small intestine after 24 h and 48 h gavage with apple snail egg PVF. **A.** Small intestine schemes highlighting parameters determined for morphometry. Left image: Section area (SA) defined as the region of interest (ROI) within the green line that includes mucosa and lumen space. Middle image: Tissue area (TA) shown in red within the ROI including only mucosa. Right image: ROI (in green) delimiting the lumen area. **B.** Changes in relative tissue area (TA). **C.** Changes in the lumen space ratio (lumen area/SA). 24 h after ingestion of *P. canaliculata* PVF, an increment in TA as well as a decrease in lumen space when compared to control group was observed. Note that these changes are partially reverted 48h after treatment where TA and lumen space begin to return to control levels. **D.** Absorptive surface (perimeter/SA) displays no changes regardless the time assayed after treatment. **E.** Typical small intestine of controls with a large lumen space and long and thin villi (arrow indicates lymphatic capillaries). **F.** Representative 24 h post-treatment section displaying a reduced lumen space with wider villi. Lymphatic capillaries (arrow) are reduced due to an increased connective tissue. **G.** Small intestine 48 h post-gavage showing villi gradually returning to control levels though its chorion still shows reduced lymphatic capillaries due to thickening of the connective tissue. Scale bar 100 μm.

**Fig 2.**
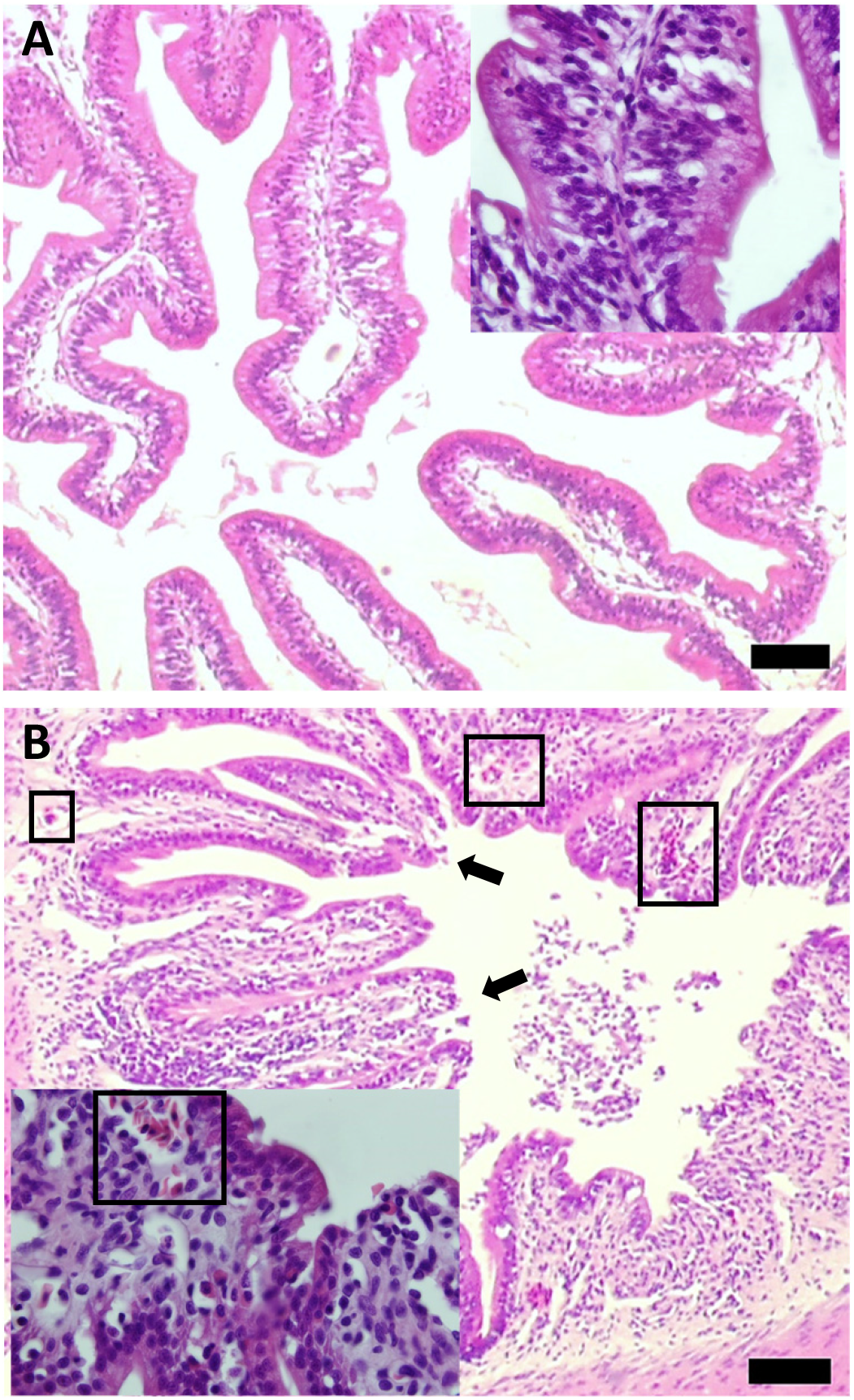
Bullfrog small intestine hemorrhage induced by *Pomacea canaliculata* eggs PVF ingestion. **A.** Representative control villi section. Inset: detail of villi chorion. **B.** 24h post-treatment showing signs of weak hemorrhages (rectangles) and epithelial disorganization at the tips of villi (arrows). Inset: detail of the weak hemorrhage (rectangle). Scale bar 100 μm.

### Gavage of egg PVF induces inflammatory response in small intestine of bullfrogs

Small intestine showed an increased number of reactive cells 24h and 48h-post-gavage as compared to control group. Eosinophils were found both in the lamina propria and between enterocytes. In control frogs 0 to 2 eosinophils were counted per field (Fig. 3A) while 24h post-gavage an increase in the number of eosinophils (7-14 eosinophils per field) was observed (Fig. 3B) decreasing to 1-6 eosinophils per field at 48 h post-gavage (Fig.3C).

**Fig 3.**
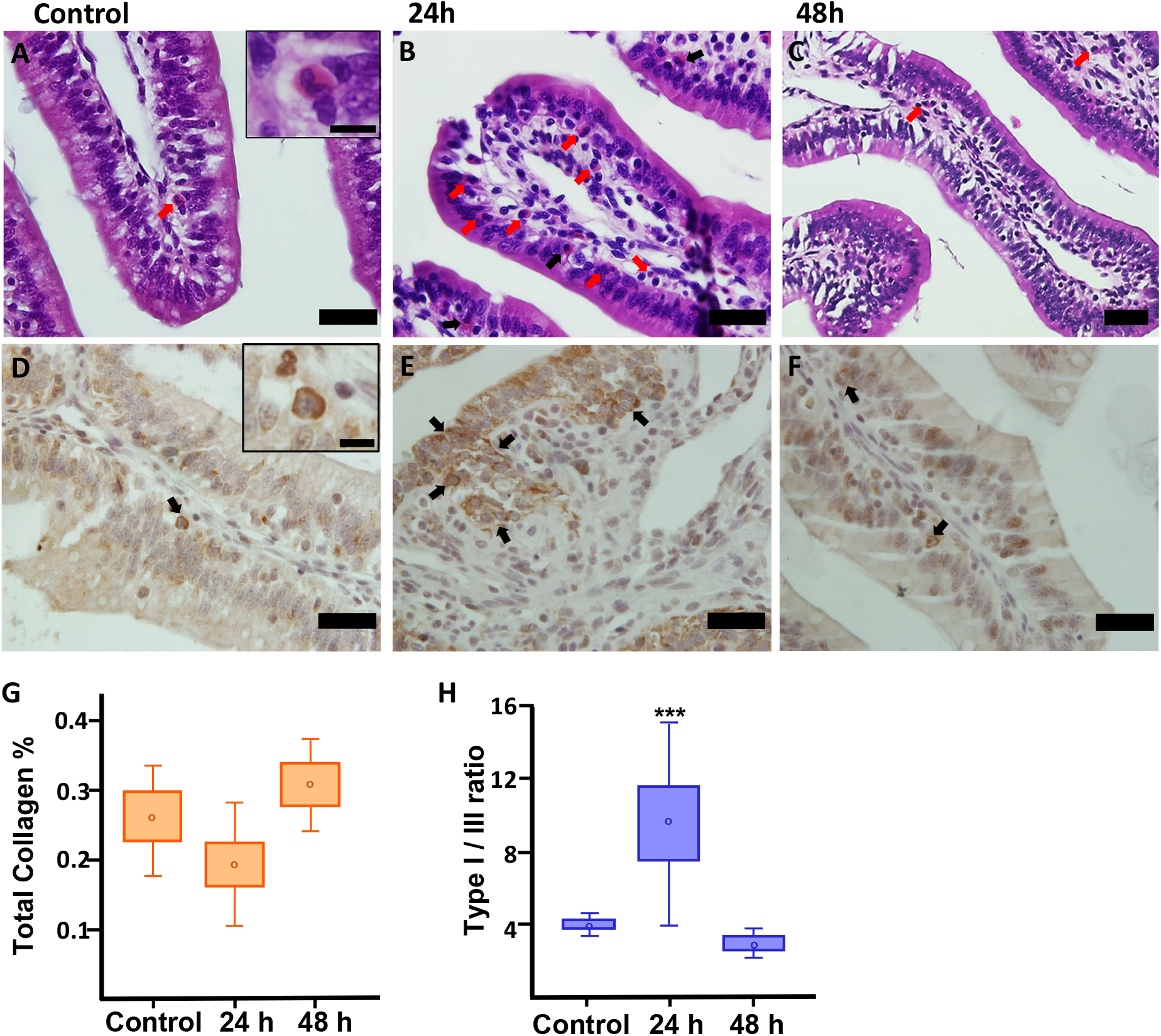
Presence of eosinophils and macrophages in bull frogs’ small intestine after gavage of *Pomacea canaliculata* eggs PVF. A-C Eosinophils (HE), D-F Macrophages (IHC). **A.** Control sections, eosinophils are marked with an arrow. Inset: eosinophil showing typical cytoplasmatic granules (scale bar 10 μm); **B.** 24h post-gavage**; C.** 48h post-gavage intestines. D-F. Representative images showing intestine macrophages (arrows) which were more prevalent 24 h after gavage with egg PVF. **D.** Control intestines showing immunolabelled macrophages with CD^68+^. Inset: detail of a macrophage (scale bar 10 μm); **E.** 24 h after-gavage; **F.** 48h after-gavage. **G**. Percentage of total collagen in the mucosa after 24 h and 48 h post gavage. H Ratio between collagen type I and III. A-F scale bar 40 μm.

The number of macrophages in the small intestine was also affected 24h post-gavage, where many immunolabelled macrophages were scattered in the lamina propria or between enterocytes (Fig. 3E). The observed number was much higher than the basal level of macrophages which were barely seen in control sections (Fig. 3D). Again, at 48 h post-gavage less labelled cells were observed than 24 h before, with a counting similar to that of the control group (Fig. 3F).

Analysis of the amount of collagen in the lamina propria showed no differences in the amount of total collagen among the three experimental groups (Fig. 3 G). However, changes in the type of collagen were observed 24 h after gavage where there was an increase in collagen I and a reduction in collagen III as compared with the other experimental groups. This resulted in an increase of collagen I: III ratio as compared with the other two groups (Fig 3 H).

### Ingestion of egg PVF induces changes in carbohydrate glycosylation pattern of enterocytes

Lectinhistochemical analysis showed differential reactivity to seven commercial lectins, particularly towards DBA, PNA and SBA (Table 1). Control sections assayed with DBA showed intense staining at the supranuclear and apical regions. Enterocytes and goblet cells were also labelled although less intensely (Fig. 4A). Twenty-four hour after gavage, enterocyte supranuclear zone was clearly negative for DBA as well as goblet cells while the apical region showed no changes as compared to controls (Fig. 4B). In animals euthanized 48 h post-gavage, staining with DBA was strong both at the supranuclear and apical regions while goblet cells display light staining (Fig. 4C). A similar weak reactivity was observed with PNA in supranuclear zone 24 h post-gavage (Fig. 4E) as compared to control group (Fig. 4D), but unlike DBA this zone stayed unstained 48 h after gavage (Fig. 4F). Goblet cells were not stained by PNA neither in control nor in treated intestines. SBA binding showed changes 24 h post-treatment where enterocytes showed stronger reactivity at the supranuclear zone (Fig. 4H) than those of control animals (Fig. 4G). After 48 h of gavage supranuclear and apical regions of enterocytes were strongly stained (Fig 4I).

**Table 1.**
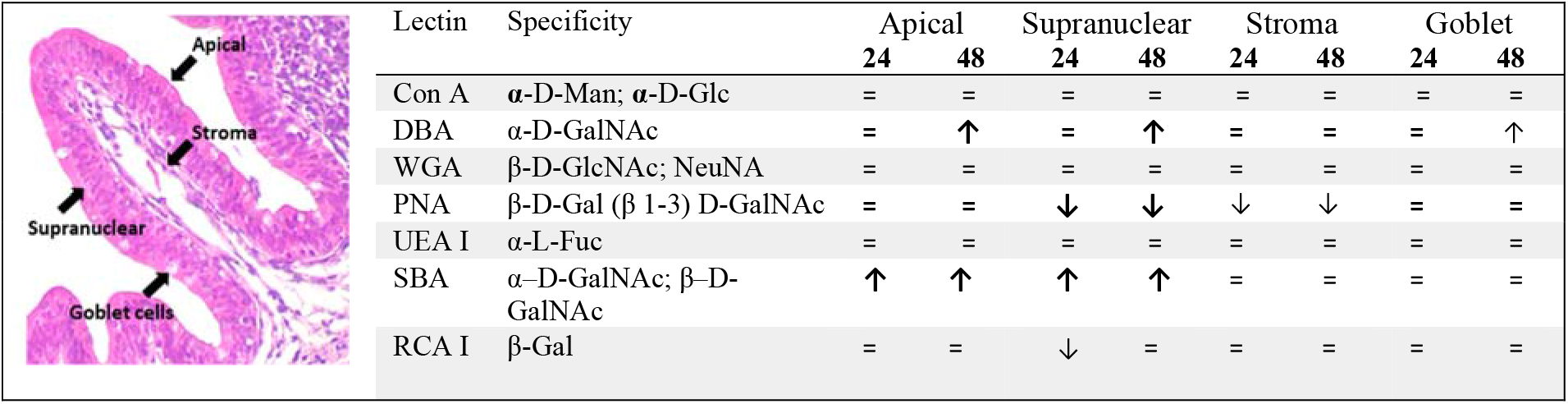
Glycosylation pattern of bullfrog small intestine villi after snail egg PVF gavage. Glycosylation changes in villi regions (see left image) were assayed by lectin histochemistry 24 h and 48 h after gavage and compared to control. No changes are symbolized with (=), and up or down arrows indicate increased or decreased reactivity, respectively. Thin arrows indicate less change (see text and Fig. 4 for details on DBA, PNA and SBA). The acronym of each lectin and their relative glycan specificity are indicated in the first two columns.

|  | Lectin | Specificity | Apical |  | Supranuclear |  | Stroma |  | Goblet |  |
| --- | --- | --- | --- | --- | --- | --- | --- | --- | --- | --- |
|  |  |  | 24 | 48 | 24 | 48 | 24 | 48 | 24 | 48 |
| | Con A | α-D-Man; α-D-Glc | = | = | = | = | = | = | = | = |
| | DBA | α-D-GalNAc | = | ↑ | = | ↑ | = | = | = | ↑ |
| | WGA | β-D-GlcNAc; NeuNA | = | = | = | = | = | = | = | = |
| | PNA | β-D-Gal ($\beta$ 1-3) D-GalNAc | = | = | ↓ | ↓ | ↓ | ↓ | = | = |
| | UEA I | α-L-Fuc | = | = | = | = | = | = | = | = |
| | SBA | α-D-GalNAc; β-D-GalNAc | ↑ | ↑ | ↑ | ↑ | = | = | = | = |
| | RCA I | β-Gal | = | = | ↓ | = | = | = | = | = |

**Figure 4.**
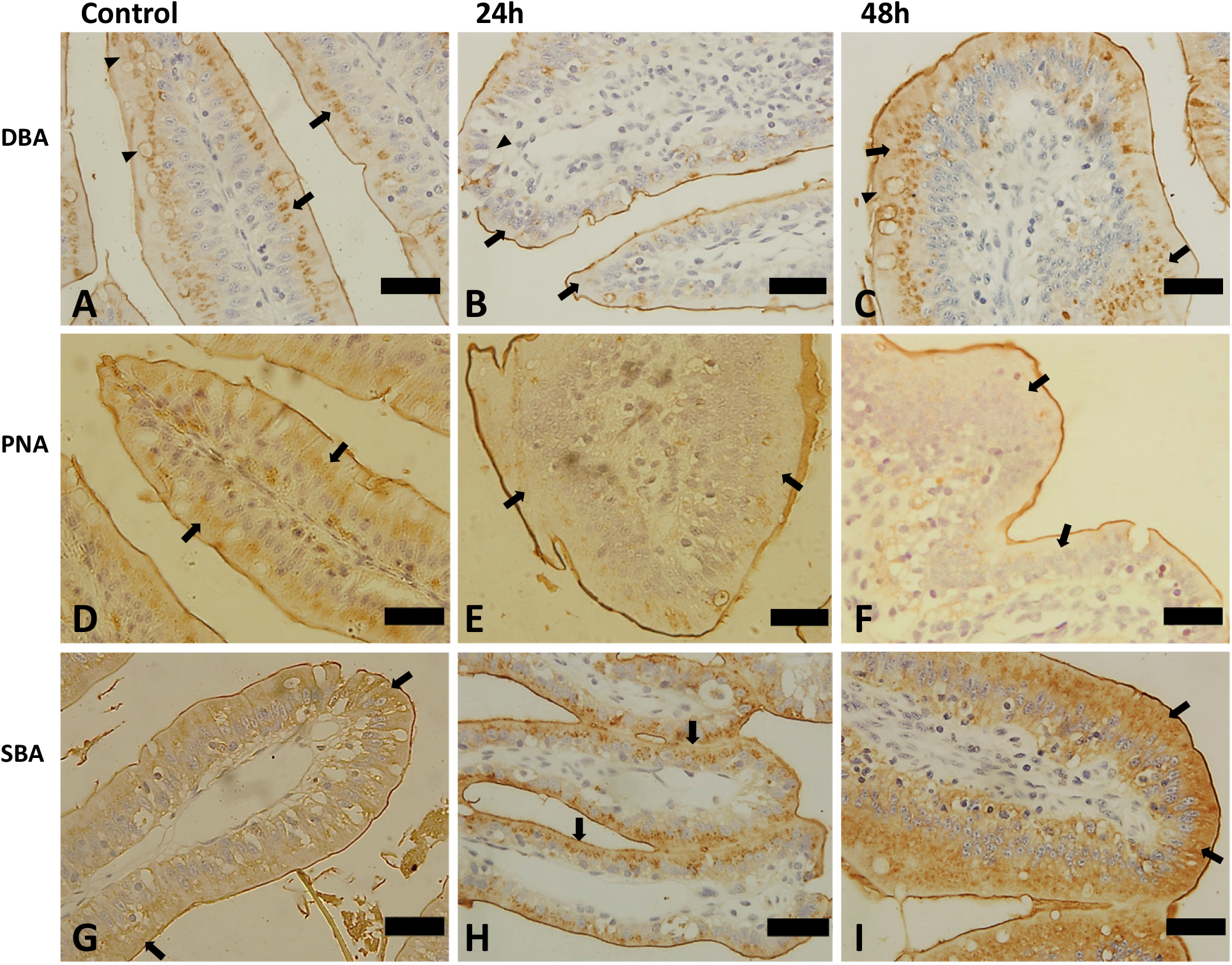
Effect of snail egg PVF ingestion on bullfrog small intestine glycosylation pattern. Lectinhistochemistry using DBA, PNA, SBA. **DBA** does not stain supranuclear zone (arrows) and Goblet cells (arrow heads) 24 h after-gavage when compared to control. These effects are reverted after 48 h post-gavage. **PNA** strongly stains supranuclear zone (arrows) in control sections although this is not seen in experimental intestines. **SBA** shows light staining in enterocytes of control animals (arrows) but a strong staining at the supranuclear zone is observed in 24 h after-gavage of experimental frogs. Staining with SBA of 48h after gavage intestines is observed not only at the supranuclear zone but also in the apical zone. Bar = 40 μm. For the complete glycosylation pattern refer to Table 1.

Interestingly, PcPV2, the enterotoxic lectin of the PVF, also binds to the intestinal epithelium, as revealed by anti-PcPV2 antibodies. The apical zone of the enterocytes facing the lumen showed clear signs of toxin immunolabelling 24 h and 48 h after gavage especially at the tip of the villi (Fig.5).

**Fig 5.**
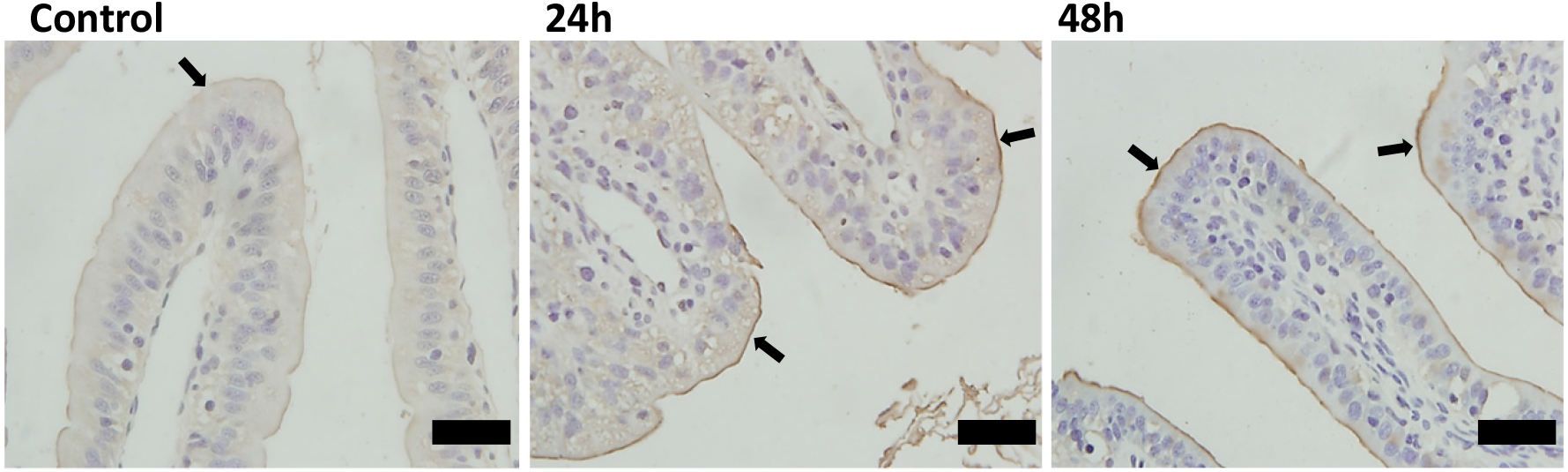
Binding of PcPV2 to bullfrog small intestine. Immunolocalization of PcPV2 at the brush border of frog enterocytes. Control group (Control) show no anti-PcPV2 antibody binding (arrow) while at 24 h post-gavage of *P. canaliculata* PVF containing the equivalent of 200μg PcPV2, a strong labelled epithelium was observed, especially ‘ at the glycocalyx of the villi (brown colour, arrows) (24 h). The same binding pattern is observed at 48 h post-gavage (brown colour, arrows) (48 h). Scale bar 40 μm.

### Intraperitoneal injection of *P. canaliculata* egg PVF is nontoxic to bullfrog

After 96 h of the i.p. injection of *P. canaliculata* PVF no mortality nor behavioural changes among juvenile bullfrogs were observed even though PVF concentration was 50 times higher than LD_50_ reported for mice. Animals were observed for any neurological associated signs, including weakness, lethargy or paralysis of rear limbs. None of these signs or any other behavioural changes were observed.

## DISCUSSION

It is expected that strong predation would select for characteristics rendering eggs inedible or more difficult to obtain. In this regard, *P. canaliculata* eggs have evolved a suite of unique biochemical defenses to avoid predation well documented in rodents. Here we extend to other taxa the knowledge of potential predators observing that ingestion of the poisonous eggs of *P. canaliculata* also adversely affect the digestive system of an amphibian.

A work performed on mammals showed that PVF have a lethal neurotoxic effect on mice within hours after i.p. injection (Heras et al. 2008). By contrast, i.p injection of PVF to bullfrogs has no lethal effect, even at a concentration 25 times higher than the concentration that would kill 100% of mice. Moreover, while sublethal doses produce rear limbs paralysis, lethargy and pain in mice none of these neurological signs were observed in bullfrogs after injection. These contrasting results between taxa suggest that the target cell or molecule for PcPV2 -the toxic lectin causing these effects on mice (Dreon et al. 2013), might be missing in anurans or, alternatively, that anurans inactivate PcPV2 in their digestive system. This last hypothesis is not supported by the fact that PcPV2 strongly binds to enterocytes *in vivo*.

Although apple snail PVF was not lethal to frogs, it severely affects their gastrointestinal tract. After 24 h of PVF gavage, intestinal morphology undergoes a dramatic change such as the increased amount of connective tissue causing an obstruction of the lymphatic vessels, and a marked epithelia disorganization at the tip of the villi. As far as we know, there is no report of similar effects induced by other potential prey on anurans upon ingestion, which limits comparisons. However, some of these morphological changes resemble those exerted by toxic anthropogenic compounds on herpetofauna (Ozelmas and Akay 1995, Çakici 2014; 2016, Çakici and Akat 2012). All changes were temporary and 48 h after gavage treated intestines resembled those of control frogs with thinner villi and the connective tissue almost reduced to the values of the non-exposed group. Moreover, some villi showed normal lymphatic vessels and nearly normal lumen space. Histopathologic effects on anuran small intestine, also observed on rats receiving these egg extracts (Dreon et al. 2014), resemble those exerted by toxic plant lectins (Vasconcelos and Oliveira 2004)that affect gut morphophysiology, at least in mammals and insects (Oliveira et al. 2004; Bardocz et al. 1995). Plant lectins, combined with digestive proteases inhibitors and kinetically stable proteins, provide an effective biochemical defense against predators (Duffey and Stout 1996; Xia et al. 2007, Van Damme 2008). Such mechanism has only been described in animals in *Pomacea* snails (Dreon et al. 2014; Pasquevich et al. 2017, Ituarte et al. 2018). *P. canaliculata* PVF protease inhibitors (Ituarte et al. 2019), lectins (Dreon et al. 2014) and enterotoxins (Giglio et al. 2020) are responsible for the changes in the gut of rodents and this may possibly be the case for anurans as well. Indeed, PcPV2 binds to the enterocyte membrane of both frogs (this study) and rats *in vivo* and to Caco-2 cells in culture (Dreon et al. 2013). The fact that the ortholog PmPV2 has enterotoxic properties on mammalian models (Giglio et al. 2020), further support that PV2 lectins from eggs would induce the histopathologic effects on frogs gut.

Enterocytes exposed to PVF also showed altered glycosylation patterns, mostly revealed by DBA, SBA and PNA. This study provides one of the few descriptions of the glycan pattern in amphibian intestine. Studies on the lectinhistochemistry of the gastrointestinal tract of anurans are scarce and on other developing stages (Kaptan et al. 2013), which limits further comparisons. Considering that glycan pattern can be affected by dietary habits, results suggest that PVF is inducing reversible changes in the surface glycosylation of the enterocytes. Altered reactivity with SBA and PNA were also observed in gut of rats fed with a diet supplemented with *P. canaliculata* PVF (Dreon et al. 2014). Taking into account that dietary lectins can induce changes in glycosylation patterns of gut epithelium (Pusztai et al. 1995) it is tempting to speculate that PVF lectins would be responsible for the observed glycosylation changes in the frog intestine.

Inflammation in frogs may be induced by a variety of physiological stressors including habitat perturbation, resource competition, parasites and pollution (Robert et al. 2014). The PVF also induced an inflammatory response 24 h after gavage with eosinophils and macrophages markedly increasing their number. Allergies may induce this cellular response though current results preclude further speculation (McGavin and Zachary 2017). Nonetheless, the response tends to revert to normal by 48 h post-gavage. This cellular reaction and the associated sudden increase of connective tissue may also be related to the enlargement of the villi and lymphatic vessel compression. Inflammatory process was not observed in rats fed with PVF under similar experimental conditions (Dreon et al. 2014; Dreon et al. 2013). Remarkably a few studies on the effect of pesticides on the digestive tract of frogs also report infiltration of eosinophils and other reactive cells (Cengiz and Unlu 2006; Velmurugan et al. 2007, Çakici 2014).

As a whole, our results indicate that 48 h after the ingestion of the PVF bullfrogs adapt to the exposure to poisonous apple snail eggs extracts. Similar temporary effects were observed in rats ingesting the same *Pomacea* egg extracts, and in rats and pigs ingesting plant lectins. It is known that the anatomy and function of the digestive tract of many vertebrate species are flexible, and can change in response to variations in environmental conditions (Karasov and Douglas 2013). Considering digestive flexibility (or plasticity) as the ability to adaptively modulate the physiology of the gut to digest different food types (Brzęk et al. 2011) we can suggest that bullfrogs have a fast-adaptive response. This is possible because the small intestine has the shortest turnover rate of all tissues in the body and, at least in mammals, it takes only 3 days to cover the entire surface with new cells. Likewise, studies on gut plasticity in amphibian, mostly performed in tadpoles (Naya et al. 2005; Ruthsatz et al. 2019), show that anurans have adaptive mechanisms that modify the activity of gastrointestinal tract (Cramp and Franklin 2005; Seliverstova and Prutskova 2012). Changes in the relative amount of collagen fibers may be related to gut reaction after ingestion of snail PVF and associated with a remodeling process although more work is needed to fully understand this issue. Our findings are consistent with the notion that bullfrogs have an adaptive response towards toxic eggs ingestion that includes a fast remodeling of the gut. To the best of our knowledge, there is no report for such response mechanism towards a potential prey in amphibians. The recently sequenced genomes of both species, the American bullfrog (Hammond et al. 2017) and *P. canaliculata* (Sun et al. 2019) provide an unprecedent model to study predator-prey interactions of two invasive aquatic organisms at a molecular level, and may suggest interesting directions for future research.

## DECLARATIONS

### Ethics Statement

Animal studies were performed in accordance with the Guide for the Care and Use of Laboratory Animals (2010) and Frogs and toads as experimental animals (Tyler 2009).

### Consent for publication

All authors have given their consent for publication.

### Availability of data and material

Data generated or analysed during this study are included in this published article and its supplementary information files.

### Authors’ contributions

TRB MSD HH conceived and designed the experiments. TBR PEF ELP MSD performed the experiments. TBR PEF ELP MSD HH analysed the data. HH contributed reagents/materials/analysis tools. TBR PEF ELP MSD HH wrote the paper.

## Acknowledgments

We are grateful to Natalia Scelcio and Guadalupe Guidi for their help with the histological work. This study was supported by a Grant from Agencia Nacional de Promoción Científica y Técnica, ANPCyT (PICT No. 2014 0850) to HH. TRB, ELP and HH are researchers of Carrera del Investigador, CONICET, Argentina.

